# Bridging Cotyledon Pathology and Perfusion in Healthy Primate Pregnancy

**DOI:** 10.64898/2026.05.18.726079

**Authors:** Logan T. Keding, Ruo-Yu Liu, Taylor J. Keding, Jessica Vazquez, Crystal G. Bockoven, Dinesh M. Shah, Thaddeus G. Golos, Oliver Wieben, Aleksandar K. Stanic

**Affiliations:** Department of Obstetrics and Gynecology, University of Wisconsin-Madison, 202 South Park St, Madison, WI, 53715 USA; Wisconsin National Primate Research Center, University of Wisconsin-Madison, 1223 Capitol Ct, Madison, WI, 53715 USA; Department of Medical Physics, University of Wisconsin-Madison, 1111 Highland Ave, Madison, WI, 53705 USA; Department of Psychology, Yale University, 100 College St, New Haven, CT, 06510 USA; Child Study Center, Yale University, 230 South Frontage Road, New Haven, CT 06520 USA; Department of Pathology and Laboratory Medicine, University of Wisconsin-Madison, 1685 Highland Ave, Madison, WI, 53705 USA; Department of Comparative Biosciences, University of Wisconsin-Madison, 2015 Linden Dr, Madison, WI, 53706, USA; Department of Radiology, University of Wisconsin-Madison, 600 Highland Ave, Madison, WI, 53792 USA

## Abstract

**Introduction:** Healthy and diseased placentae alike often display some degree of pathology. However, quantitative techniques to characterize common pathologies and their relationship to local maternal hemodynamics in healthy primate placentae are currently limited.

**Methods:** Placentae from seven rhesus macaques were imaged by MRI at three time points across mid-to late-gestation, to quantify placental blood volume, flow, and perfusion from maternal spiral arteries across pregnancy. Near term, we collected placental cotyledons, digitized hematoxylin/eosin-stained slides, then segmented and annotated sub-tissues and major pathologies (intervillous gaps, fibrin deposition, villous agglutination, inflammatory agglutination, and stromal mineralization) within each cotyledon. Individual pathologies were assessed in relation to each other and MRI perfusion metrics, in a cotyledon-specific manner. Parallel analyses were performed to investigate both basic (Spearman correlation) and animal variance-negated (dimensionality-reduction) relationships.

**Results:** Cotyledons with increased stromal mineralization demonstrated low blood perfusion across pregnancy, alongside significant compensatory changes. Mineralization was further associated with decreased fetal weight, across all sub-tissues. Dimensionality reduction revealed maternal vascular malperfusion-associated pathologies as the largest contributor to dataset variance. Additionally, pathologies commonly associated with healthy placental function demonstrated low cotyledon blood flow and volume at all timepoints, with no evidence of compensatory changes across gestation.

**Conclusions:** Comprehensive digital annotation revealed several relationships connecting pathology and maternal blood perfusion in the healthy primate pregnancy, at the smallest functional unit of the placenta. This methodological framework embeds pathologist-refined morphological expertise into a quantitative, spatially resolved format that can ground, rather than be replaced by, unsupervised computational approaches to placental analysis.

## INTRODUCTION

Major adverse pregnancy outcomes (APOs) are commonly attributed to placental insufficiency [1], often determined through analysis of fetoplacental tissues. Rarely, lesions indicative of insufficiency can be identified macroscopically [2], though they are more commonly observed at the microscopic level [3]. These microscopic pathologies are determined through visual assessment by placental pathologists and compose the major pathological meta-categories: maternal vascular malperfusion, fetal vascular malperfusion, and chronic inflammation [2,4–6]. Pathologists relay their findings in written reports, using qualitative or semi-quantitative description of injury determined through visual estimation [4]. Although these methods have helped standardize placental injury characteristics clinically, they have not provided the resolution required to link pathological manifestation with pregnancy outcome [2].

There is a basic consensus that placental pathologies lead to adverse pregnancy outcome, however, the threshold of injury that results in pregnancy disease states remains undetermined [2]. For example, 50% of placentae that meet maternal vascular malperfusion criteria and 19% of placentae exhibiting chronic inflammation result in healthy pregnancy outcome [7,8]. Additionally, 20% of all pregnancies that display obstetric complications show healthy placental findings [9]. These seemingly contradictory observations likely result from a combination of current placental analysis limitations, such as: 1) poor sampling: sparse and selective sampling of macroscopic pathologies leading to oversampling of easily identifiable lesions; 2) poor resolution: visual estimation and qualitative descriptors like “moderate” or “severe”, resulting in diminished resolution/translatability to other strictly quantitative forms of biological data; and 3) unrecorded analyses: visual pathological assessment allows scant opportunity for secondary review and revision of primary analysis.

In humans, the placenta is divided into vascular functional units termed cotyledons [10], which are perfused by one, or multiple maternal spiral arteries [11,12]. These cotyledons are comprised of specialized regions: the chorionic plate, intervillous space, placental villi, and trophoblastic shell [13], in addition to the decidua basalis, a tissue of maternal origin [14]. The rhesus macaque placenta shares these attributes with the human, along with a discoid placental structure, hemochorial villous placental architecture, and near identical maternal hemodynamics [12,13,15]. These characteristics, combined with the ability for experimental manipulation, make the rhesus macaque an ideal candidate for placental study and translatability to the human [16].

Utilizing the macaque, we demonstrate a novel quantitative, spatially resolved placental pathology analysis approach. Through extensive digital slide segmentation and annotation, we successfully mapped major sub-tissues and pathologies within healthy macaque cotyledons. We then leveraged this methodology in conjunction with established dynamic contrast enhanced (DCE) magnetic enhanced imaging (MRI) techniques [17] towards understanding the local connection between placental pathology and maternal blood delivery in the primate (Figure 1). Through these efforts, we report relationships that connect inter-pathological manifestation, pathology with compensatory perfusion, and pathology with diminished fetal growth; all from placentae of primate pregnancy with healthy/normal outcomes. Importantly, this new paradigm of digital placental pathology annotation can be well utilized within the current pregnancy research context; where spatially resolved, quantitative data analysis has become increasingly pivotal.

**Figure 1.**
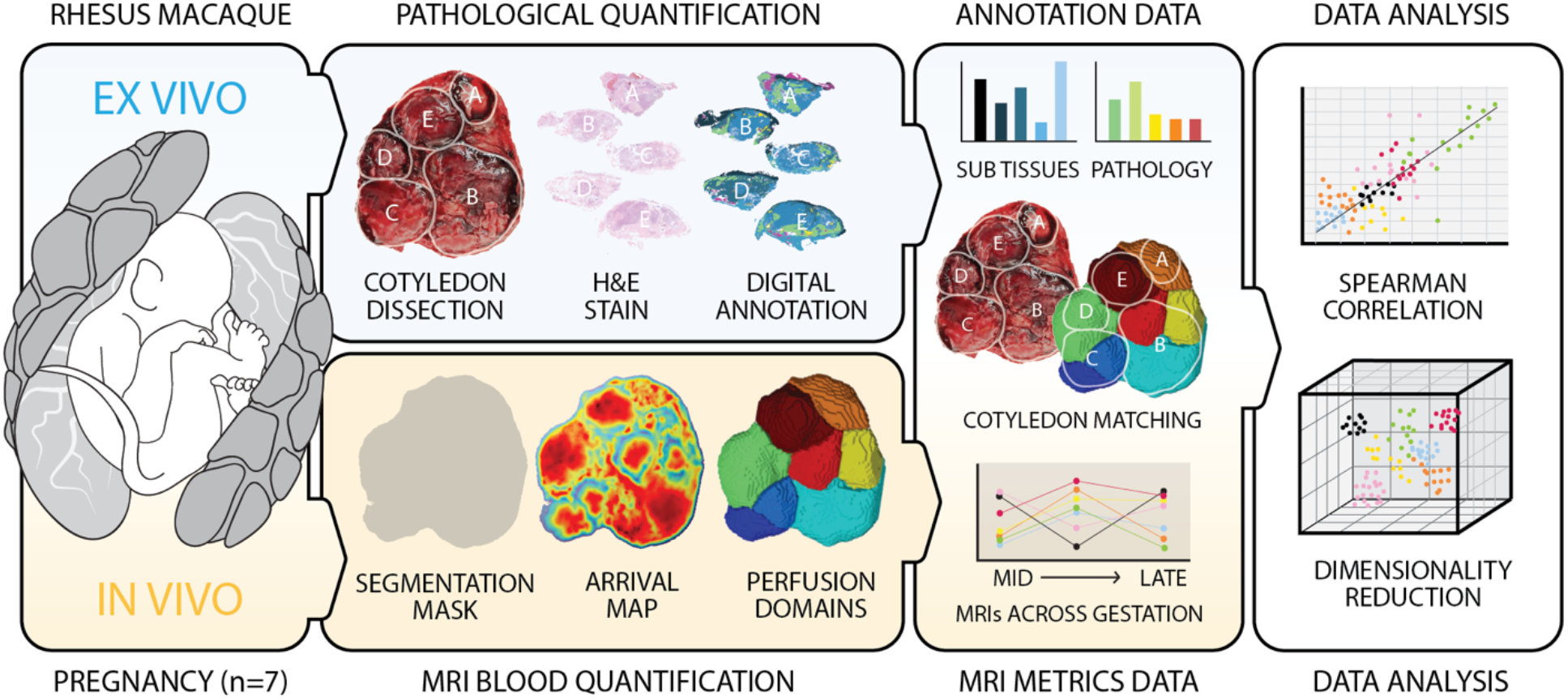
Experimental Design Overview. Seven pregnant rhesus macaques received *in vivo* constrast-enhanced MRI at three timepoints from mid-to late-gestation alongside *ex vivo* placental pathology analysis at term via digital annotation. Afterwards, annotation and MRI datasets were paired by cotyledon, then analyzed by Spearman correlation and dimensionality reduction for significant relationships.

## METHODOLOGY

### Macaque Injections, MRI, and Placental Preparation

All experimental procedures were performed in accordance with the NIH Guide for the Care and Use of Laboratory Animals and with the approval of the University of Wisconsin College of Letters and Sciences and Vice Chancellor Office for Research and Graduate Education Institutional Animal Care and Use Committee, protocol G006209.

Seven saline-injected pregnant rhesus macaques were recruited from multiple WNPRC studies [17,18] to serve as healthy/normal primate pregnancies for this study. Three macaques were injected with 1 mL of saline into the amniotic sac at gestational day (GD) ~55 [17], while four additional macaques received a 0.5 mL saline injection to the anterior disc at GD ~100 [18]. The macaques underwent MRI procedures on clinical 3T scanners at three separate timepoints to track local changes across gestation: mid- (GD 65-95), mid/late- (GD 100-115), and late-gestation (GD 137-147). Each exam included a DCE-MRI scan using ferumoxytol (Feraheme, AMAG Pharmaceuticals), a pregnancy safe contrast agent used to assess maternal blood dynamics, as described previously [18–20]. Data were processed using an established workflow [17] to obtain measures of cotyledon volume (mL), blood flow (mL/min), and perfusion (blood flow [mL/min]/volume [mL]) for each perfusion domain. The perfusion domains were identified by DCE MRI through an automated analysis based on contrast arrival times adopted from our groups’ previous work [21].

Pregnant macaques underwent cesarean section to excise placental and fetal tissues at GD ~155, short of full gestation (GD ~167) [22] to avoid early labor. Placental cotyledons were determined through visual and textural assessment, then dissected along placental septae (primary disc n = 50, secondary disc n = 30). Once isolated, full-thickness sections of the central cotyledonary mass were obtained. Cotyledon center cuts were paraformaldehyde-fixed, paraffin-embedded, sectioned, stained with H&E, and scanned at 4x magnification as previously described [17,18]. The MRI-derived perfusion domains identified at each gestational timepoint were then matched to the cotyledons identified in the postpartum placentae based on relative location, size, and boundaries using photographs and matching MRI reformats (Figure 1).

### Cotyledon Annotation Overview

All cotyledon sub-tissue and pathology annotation categories were developed and performed by LTK with counsel from CGB, a board-certified placental pathologist. Final annotation quality was successfully verified by CGB through assessment of 10 randomly selected cotyledon images. Cotyledon center cut images were assigned random numeric identifiers, resulting in annotation blind to animal and placental disc origin. Annotations were performed in open-source GIMP (GNU Image Manipulation Program) version 2.10.38. Each sub-tissue and pathology comprised a discrete layer, of which pixels could be selected, enumerated, and assessed for overlap with other layers. Free Select, Select by Color, and Fuzzy Select tools were used to select pixels, while the Histogram data tool was used to quantify the number of pixels in each selection/layer. Cotyledon sub-tissue layers constituted: the chorionic plate, placental villi, intervillous space, trophoblastic shell, and decidua basalis (Figure 2, Supplemental Figure 1). All sub-tissues were identified through tissue staining and structure, representing non-overlapping areas (Supplemental Table 1). The distinct origin and immune environment of the decidua basalis, in combination with the sparsity of the tissue at term, led to our exclusion of this tissue from downstream analyses.

**Figure 2.**
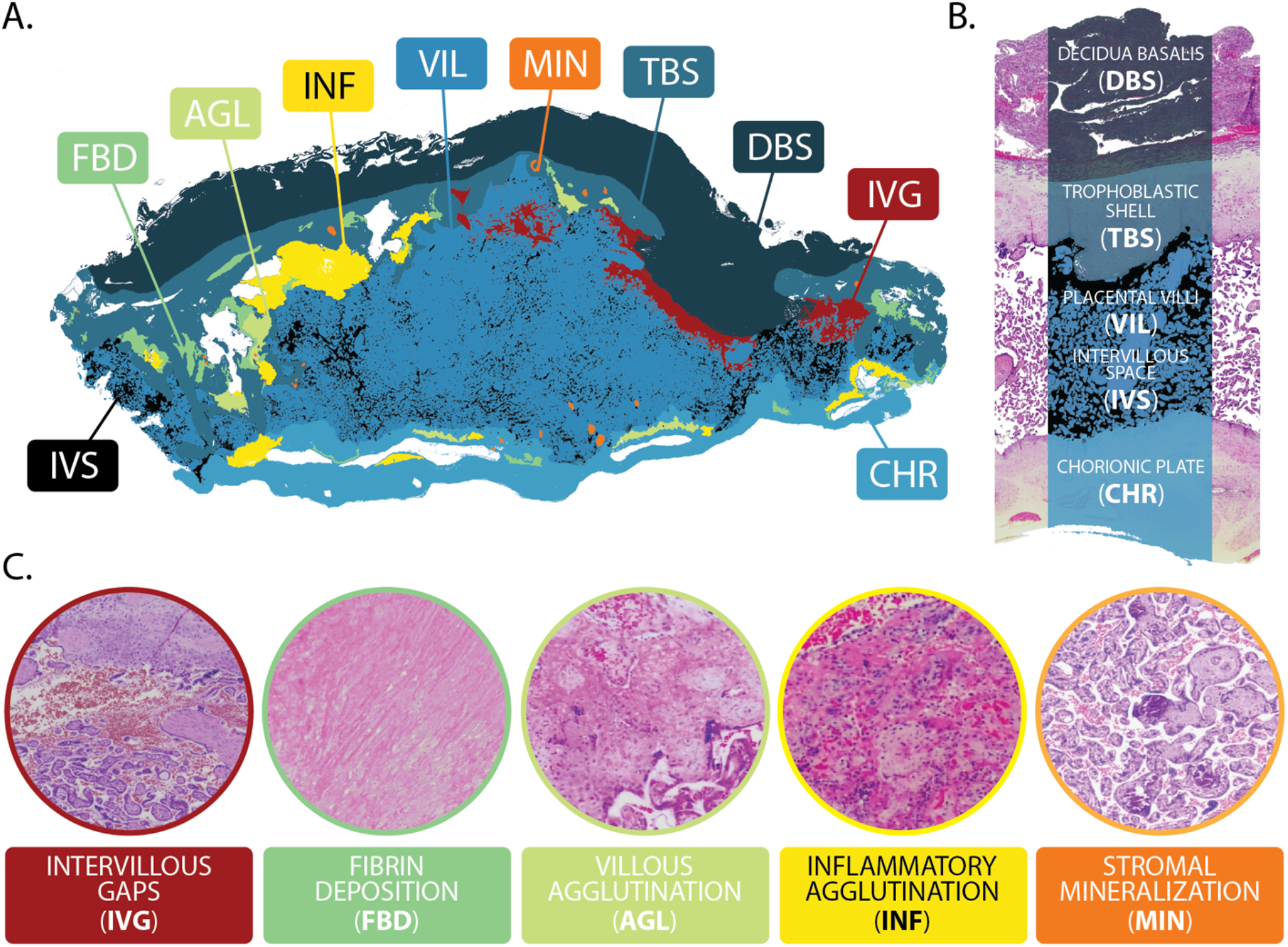
Cotyledon Sub-tissue and Pathological Annotations. A) A representative annotated placental cotyledon, displaying all potential sub-tissues and pathologies. B) A representative overlay of sub-tissue annotations: decidua basalis (DBS), trophoblastic shell (TBS), placental villi (VIL), intervillous space (IVS), and chorionic plate (CHR). C) Representative, color-coded examples of pathology annotations: intervillous gaps (IVG), fibrin deposition (FBD), villous agglutination (AGL), inflammatory agglutination (INF), and stromal mineralization (MIN). All tissues were H&E stained and scanned at 4x magnification.

Pathological annotations consisted of 1) gaps within the intervillous space, 2) fibrin deposition, 3) villous agglutination, 4) inflammatory villous agglutination, and 5) stromal mineralization (Figure 2, Supplemental Figure 1). Intervillous gaps were defined as significant gaps or blood pooling within the intervillous space along the trophoblastic shell and/or central parenchyma (Figure 2, Supplemental Figure 2). Fibrin deposition was identified as heavily-stained eosinophilic material (Figure 2, Supplemental Figure 3), displaying varying degrees of nucleation [24]. Villous agglutination was defined as architectural damage of the villi, combined with syncytial necrosis and inter/perivillous fibrin deposition (Figure 2, Supplemental Figure 4). Inflammatory villous agglutination was defined the same as villous agglutination, with the addition of heavy leukocytic infiltrate (Figure 2, Supplemental Figure 5). Stromal mineralization was identified by regions of dark, hematoxylin-rich stain within villous stroma (Figure 2, Supplemental Figure 6). Additional inclusion criteria, exclusion criteria, and associated historical pathological terminology for pathological annotations can be found in Supplemental Table 2.

To account for variation in cotyledon size, sub-tissue areas were normalized by total cotyledon area. Similarly, pathological pixel areas were reported as a proportion of sub-tissue area (i.e. chorionic plate mineralization pixels/chorionic plate pixels). GraphPad Prism (version 10.4.1) was used to calculate non-parametric Spearman tests for all correlation analyses, to account for prevalent non-normality and zero-inflation of our dataset.

### Dimensionality Reduction of Pathology Markers

As genetic composition and environmental factors both contribute to the manifestation of placental pathology [28], we used dimensionality reduction to limit animal-specific contributions to pathological relationships using partial distance-based Redundancy Analysis (dbRDA) with the vegan package (vegan package 2.7-2) in R [29–31]. The final dataset comprised nested observations, defined by “cotyledon” within “animal.” First, a dissimilarity matrix was calculated using the Bray-Curtis distance; chosen for its robust handling of zero-inflated, proportion-level data. To isolate within-animal variation and control for baseline differences between animals, dbRDA was implemented as an intercept-only model conditioned on the animal (Supplemental Figure 7):

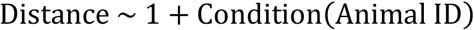

Principal coordinate analysis (PCoA) eigenvectors of the dissimilarity matrix were calculated after orthogonalizing against animal and partitioning out animal-related variance. Consequently, the resulting unconstrained ordination axes (i.e. components) reflected the intra-animal heterogeneity of pathology markers across cotyledons.

Significant pathological principal components were identified using a permutation-based Mantel test (n=1000 permutations). For each component retrieved from the dbRDA, the Euclidean distance matrix of the component scores was calculated and correlated with the original Bray-Curtis dissimilarity matrix. Components were retained if they met the following criteria: a Mantel test p < 0.05 (chance criteria) and Variance Explained/Cumulative Variance Explained < 0.05 (noise criteria) [32]. To aid in the biological interpretation, we calculated the Spearman rank correlation coefficients (ρ) between the original pathology markers and the component scores. The statistical significance of these loadings was assessed using additional permutation testing (n=1000 permutations) where the pathology marker vectors were randomly shuffled to generate a null distribution of correlations. Markers with a permutation *p* < 0.05 were considered significant drivers of the respective pathology component (Supplemental Figure 8). All analysis software is available at https://github.com/staniclab/placenta-pathology-dbRDA.

To determine if the identified pathology components predicted MRI metrics across gestation, we utilized linear mixed-effects models via the *lme4* (v 2.0-1) and lmerTest (version 3.2-1) packages in R. We analyzed three perfusion metrics with independent models at the cotyledon level: volume, blood flow, and perfusion; at three unique timepoints. A random intercept for animal ID was included to account for baseline animal-related differences in blood perfusion. Model selection was performed using backward elimination: interaction terms with p > 0.05 were removed, and the models were refit with main effects and surviving interactions only. Final model parameters were standardized, and *p*-values for fixed effects were corrected for multiple comparisons using the false discovery rate (FDR) method [33]. All relationships were visualized using residual plots to isolate the specific effects of interest.

## RESULTS

### Cotyledon Descriptive Statistics

To quantify basic placental features of this cohort, sub-tissue and pathological annotations were completed for all cotyledons, across all placentae. The average contributions of cotyledon sub-tissues and individual pathologies in relation to total cotyledon and total pathological area are presented in Figure 3. As is also observed in human placentae, the villous region encapsulated most of the total cotyledon area, followed by the trophoblastic shell (Figure 3) a more prominent feature of macaque placentae when compared to humans [34]. The most common pathology by pixel area was fibrin deposition, followed by intervillous gaps and stromal mineralization, while inflammatory and non-inflammatory villous agglutination constituted the smallest proportion of total pathology (Figure 3). Within the cotyledon, most injury manifested near the trophoblastic shell (Figure 3), in agreement with observations in human placentae [2]. Both inflammatory and non-inflammatory villous agglutination most often occurred along the trophoblastic shell and chorionic plate borders, whereas fibrin deposition and stromal mineralization most often occurred within the villous region (Figure 3).

**Figure 3.**
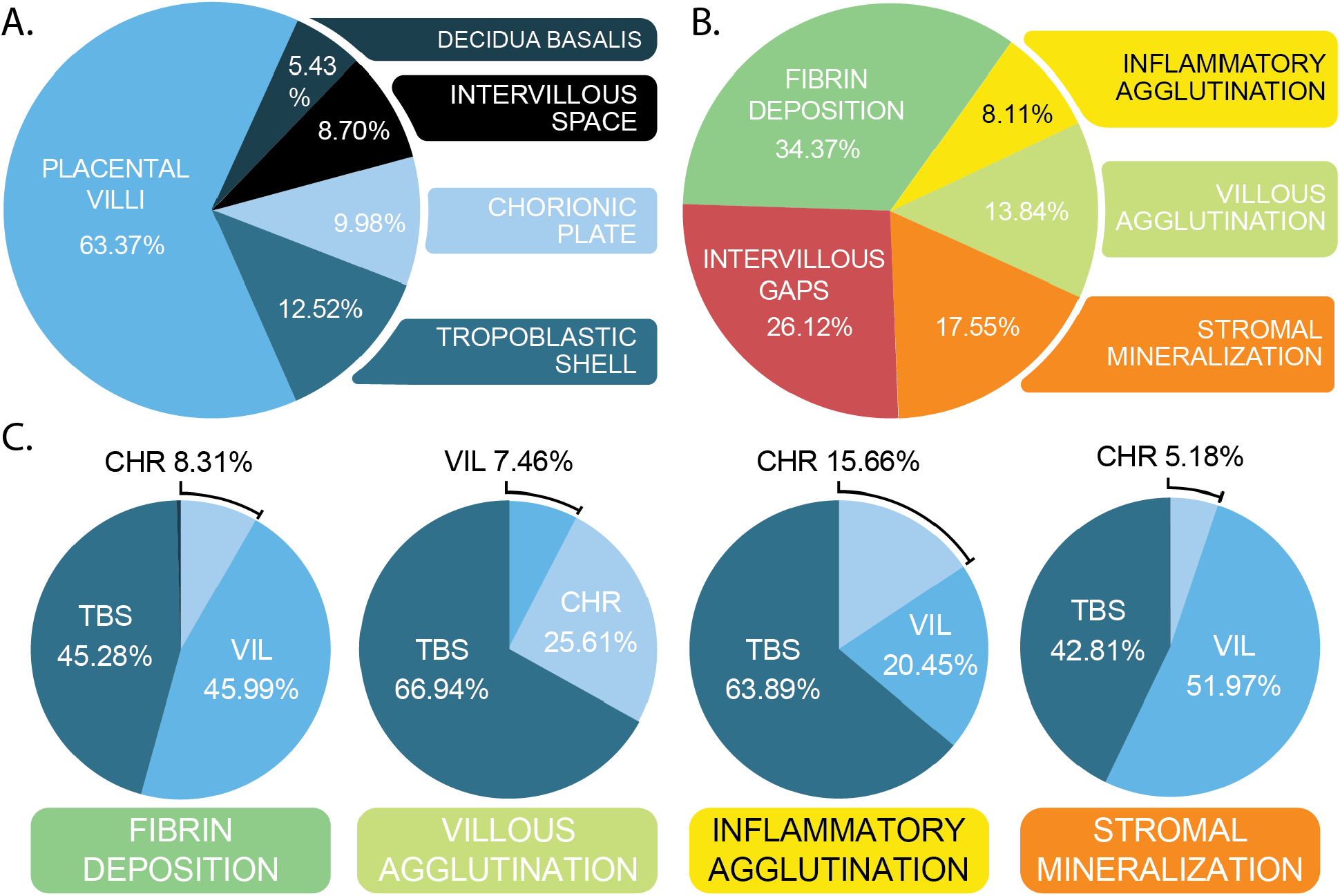
Descriptive Statistics of Cotyledon Sub-Tissues and Pathologies. A) Percent contribution of cotyledon sub-tissues to total cotyledon tissue area by pixel area. B) Percent contribution of individual cotyledon pathologies to total cotyledon pathology by pixel area. C) Percent expression of individual pathologies by cotyledon sub-tissue. Decidua basalis not represented due to negligable contribution (<0.05%).

### Cotyledon Pathology Relationships

Placental lesions associated with adverse pregnancy outcome require a substantial parenchymal coverage to warrant diagnosis [4,5] and demonstrate a high degree of correlation in pathological pregnancy [2]; however, less is known about their manifestation in healthy pregnancy. Therefore, we sought to investigate the relationship between sub-threshold quantities of pathology in healthy macaque cotyledons (Figure 4A and B).

**Figure 4.**
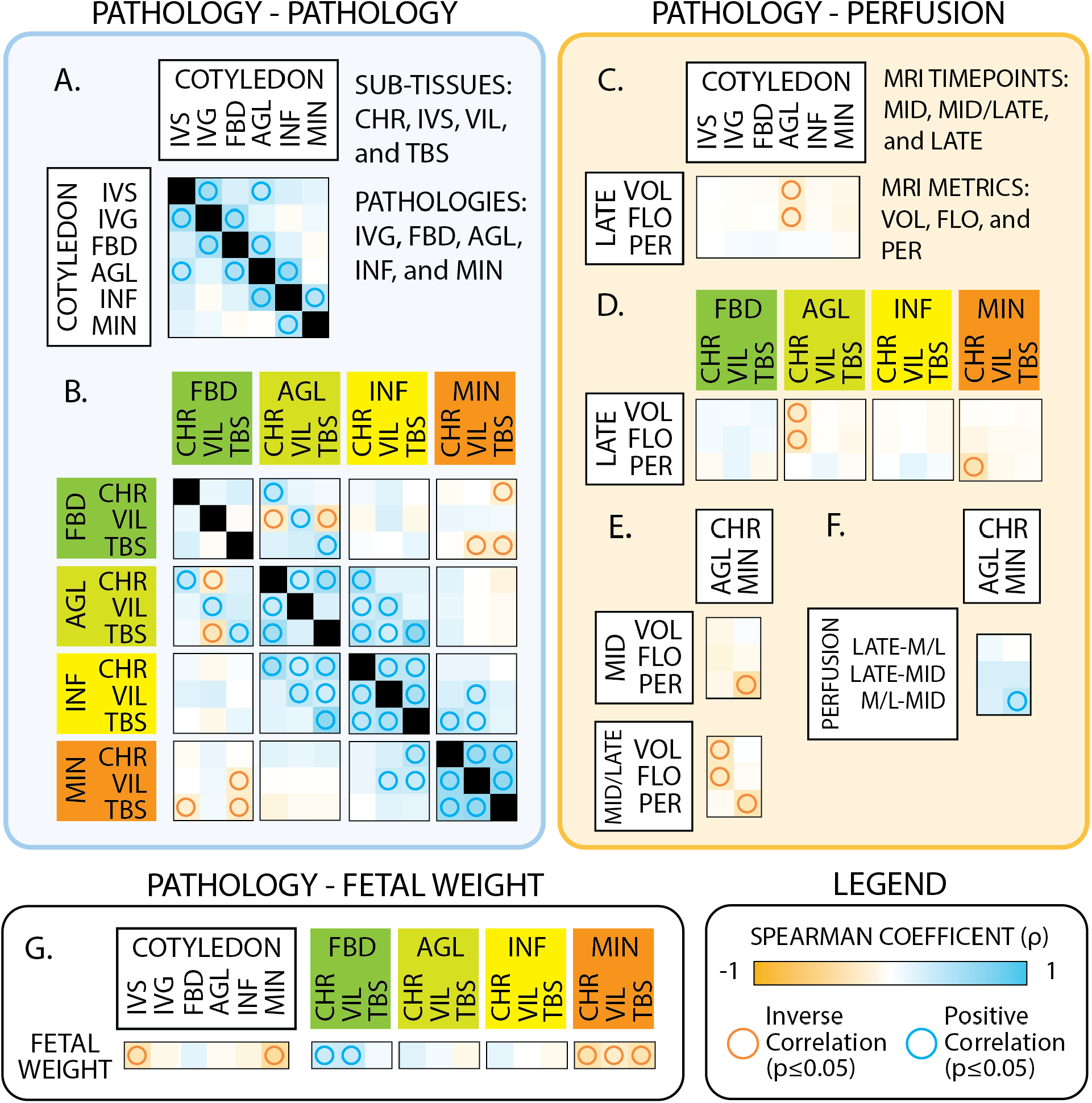
Connecting Placental Pathology, Perfusion, and Fetal Weight by Spearman Correlation. Individual pathological correlations were first assessed at cotyledon (A) and sub-tissue (B) levels. Next, individual pathologies were assessed in relation to late-gestation MRI metrics (volume [mL,VOL], blood flow [mL/min, FLO], and perfusion [mL/min/mL, PER] at cotyledon (C) and sub-tissue (D) levels. AGL and MIN pathologies were further investigated in relation to MRI metrics at earlier mid- and mid/late (M/L)-gestation timepoints (E). Perfusion was further assessed in relation it’s change across mid-, mid/late- (M/L), and late-gestation timepoints (F). Last, individual pathologies were assessed in relation to term fetal weight at cotyledon and sub-tissue levels (G).

Total intervillous space, intervillous gaps, fibrin deposition, and villous agglutination demonstrated positive associations across cotyledons (Figure 4A), all histological features associated with maternal vascular malperfusion (MVM) when significantly elevated. Inflammatory and non-inflammatory villous agglutination showed a strong positive relationship at the cotyledon and sub-tissue levels (Figures 4A and B), related pathological features shown to manifest in tandem clinically [35]. Notably, stromal mineralization shared a single positive relationship with inflammatory villous agglutination at the cotyledon level and sub-tissue level (Figures 4A and B). Except for fibrin deposition, each pathology demonstrated a strong, positive association with its own occurrence across sub-tissues (Figure 4B).

### Stromal Mineralization, Maternal Perfusion, and Fetal Weight

Next, we sought to investigate the relationship between maternal blood perfusion with annotated vascular malperfusion- and chronic inflammation-associated lesions. Non-inflammatory villous agglutination was the sole pathology significantly related to maternal blood dynamics at the cotyledon-level, demonstrating that agglutination increased in smaller, slower flowing cotyledons (Figure 4C). When interrogating at the sub-tissue level, we found that agglutination at the chorion most significantly contributed to this relationship (Figure 4D). In addition, stromal mineralization at the chorionic plate increased as maternal blood perfusion decreased locally (Figure 4D).

To determine if these chorionic agglutination and mineralization pathologies at term were related to altered MRI metrics across gestation, earlier timepoints (mid- and mid/late-gestation) were investigated (Figure 4E). Chorionic agglutination was inversely associated with cotyledon volume and blood flow at mid/late-gestation, while chorionic mineralization was associated with diminished perfusion at both mid and mid/late timepoints (Figure 4E). We further assessed chorionic agglutination and mineralization in relation to maternal blood metric changes between each MRI timepoint (1] mid/late-mid, 2] late-mid, and 3] late-mid/late) (Figure 4F). The sole significant finding was an increase in chorionic mineralization as perfusion increased between mid- and mid/late-gestation (Figure 4F); demonstrating a connection between compensatory changes in perfusion and increased chorionic plate stromal mineralization at term.

Finally, as significant increases in placental pathologies are associated with decreased fetal growth [36,37], we sought to quantify the relationship between cotyledon pathology and fetal weight (Figure 4G). Increased cotyledon intervillous space and stromal mineralization across all sub-tissues were the sole pathologies associated with decreased fetal weight (Figure 4G). As fetal weight is an animal-level endpoint rather than a cotyledon-level measurement, the Spearman correlations presented here are descriptive, rather than confirmatory.

### Component Analysis of Pathological Relationships

Dimensionality reduction resulted in the generation of significant pathological relationships, termed “components”, which in total accounted for ~73% of the dataset’s total variance (Supplemental Figure 8). Components 1 and 2 encapsulated pathological features commonly associated with maternal vascular malperfusion (MVM) (Figure 5). Component 1 explained 22.8% of total variance, where increased total area and gaps within the intervillous space proved most significant, followed by coagulation-associated pathologies at the maternal floor, as well as chorionic AGL (Figure 5). Increased intervillous space and gaps in the placenta are associated with accelerated villous maturation and altered local maternal blood supply [38], as is the agglutination of villi. Largely absent from this component were all FVM- and chorionic inflammatory-associated annotations (Supplemental Figure 8). Component 2 accounted for an additional 16.8% of total variance, further specifying MVM pathologies of the maternal floor (Figure 5). The separation of MVM-associated components 1 and 2 likely reveal deviations in origin of injury, maternal hemodynamics, and/or villous development in relation to malperfusion lesion manifestation.

**Figure 5.**
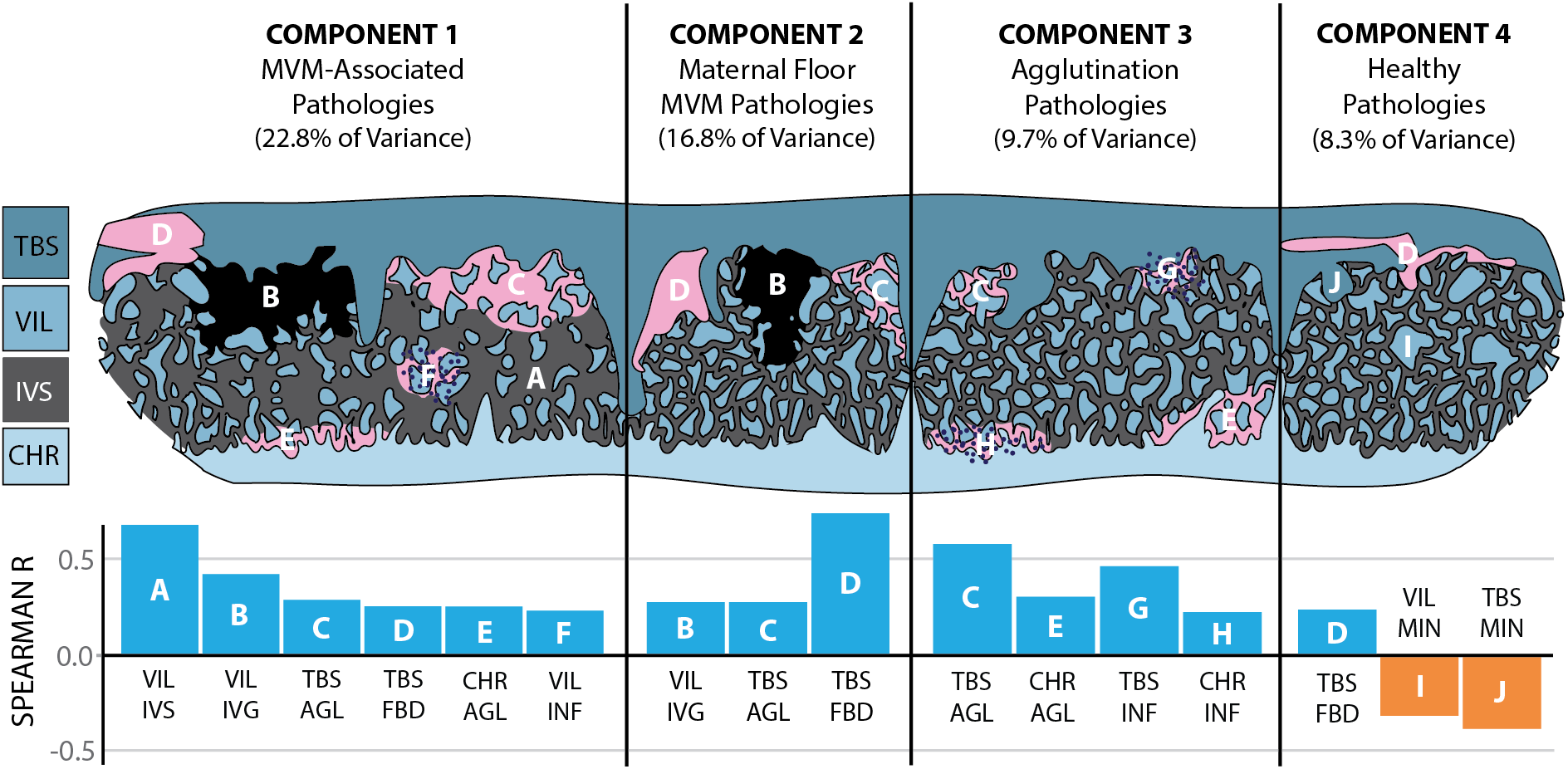
Placental Pathological Components by Partial Distance-Based Redundancy Analysis (dbRDA). The dimensionality of pathological annotations was reduced to major pathological components. Top) The top four significant components contributing to data variance with Middle) graphical representations of type and location of individual significantly contributing pathologies. Bottom) Bar graphs depicting Spearman rho values of individual type and location of pathologies significantly contributing to components 1-4. Letters (A-J) relate type and location of pathology bar graph data to graphical representations above.

Component 3 explained 9.7% of remaining variance, highlighting the co-occurrence of inflammatory and non-inflammatory villous agglutination along the periphery of the villous region (Figure 5). This component supports high co-expression of MVM and chronic inflammatory pathology and a potential common etiology in some proportion of cases, as has been noted previously [35]. Component 4 describes 8.3% of the remaining variance, highlighting the relationship between increased trophoblastic shell fibrin deposition alongside decreased trophoblastic shell and villous stromal mineralization (Figure 5). As trophoblastic shell fibrin was associated with MVM pathologies in two prior components, it is likely that this component represents physiological fibrin formation at the basal plate; hypothesized to serve a beneficial function at the placenta, as opposed to being evidence of injury [26]. As absence of significant stromal mineralization is undoubtedly also a sign of villous health at the placenta [36], we determined the variance accounted for by this fourth component represents healthy pathological relationships at the placenta.

### Healthy Pathologies are associated with Low Cotyledon Volume and Blood Flow Across Gestation

Gestational age was positively associated with cotyledon blood volume and flow, and negatively associated with perfusion (Supplemental Figure 9), as previously observed [17,39,40]. Surprisingly, we did not observe any significant relationship between MVM pathology components 1 and 2 and altered placental perfusion metrics. However, a relationship between the healthy pathological manifestation represented by Component 4 and decreased cotyledon volume and blood flow across gestation was observed (Figure 6), with the relationship to blood flow being particularly stark, remaining significant following FDR correction (Figure 6B). Simultaneously, perfusion across gestation shared no relationship with component 4, suggesting the low volume and blood flow observed did not equate to lower perfusion.

**Figure 6.**
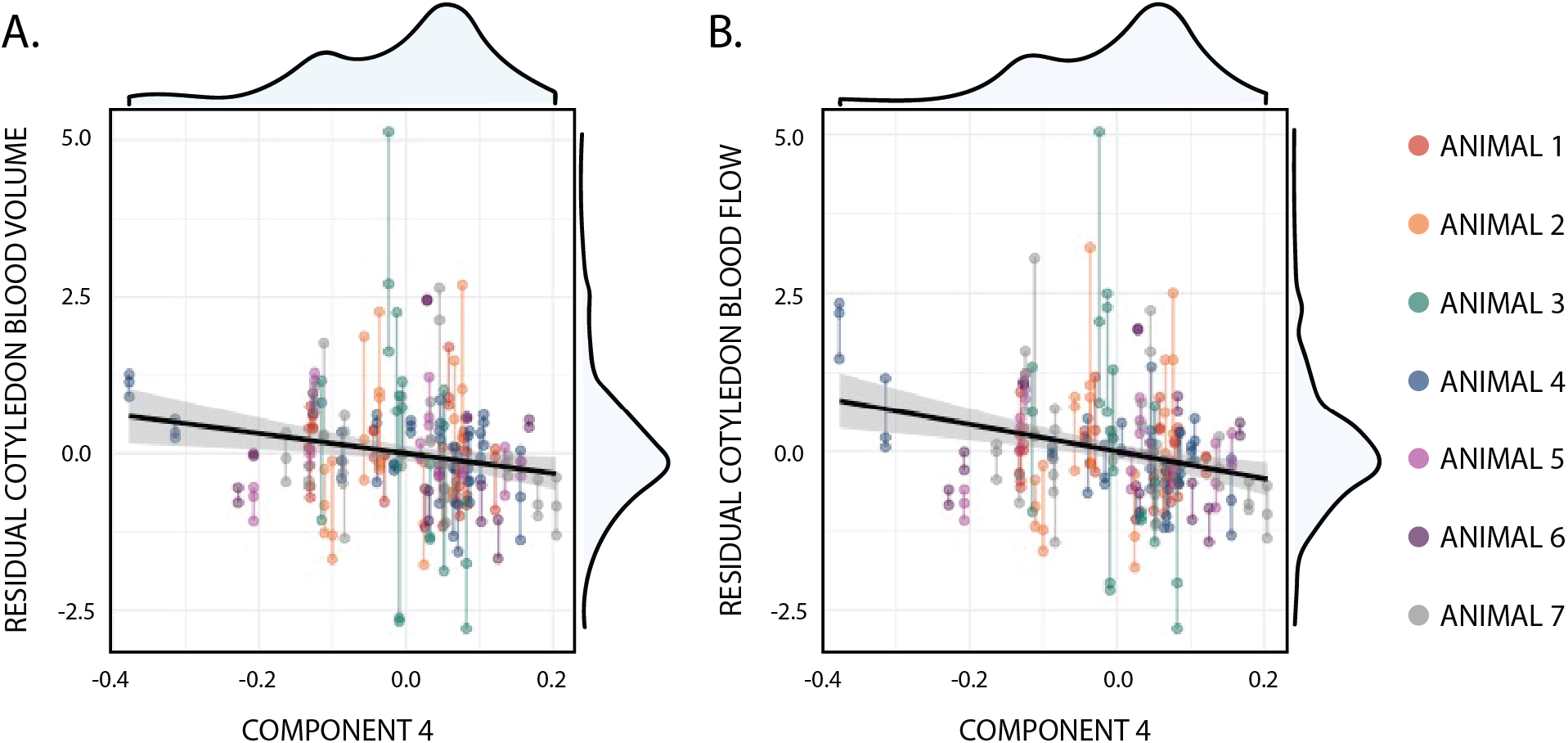
Pathology Component 4 relationship to Longitudinal Blood Volume and Flow. Dot plots depicting blood volume (A) and flow (B) measurements in relation to pathological Component 4. Individual dots represent values from mid-, mid/late-, and late-gestation, joined by a central line to connect all timepoints from the same cotyledon. Cotyledon dots and lines are associated with the rhesus macaque they were derived from by color. Both relationships were found to be significant, with blood flow vs Component 4 remaining significant following FDR correction.

## DISCUSSION

The placental injury that distinguishes healthy from adverse outcome pregnancy remains poorly resolved [2]; in part because injury characterization has relied on qualitative, whole-organ-level assessment that obscures local relationships between pathology and perfusion. MVM, FVM, and chronic inflammatory pathologies all contribute to pregnancy complications [3–5], but are also found within healthy/normal placentae [2,7,41,42]. The primary innovation presented here rests on the methodological shift to strict digital quantification, as historical analysis relies on visual estimation and qualitative descriptors that lack annotation record, quantitation, and reproducibility. Second, our study investigates the smallest functional unit of maternal placental perfusion; connecting longitudinal changes in blood dynamics measured using state-of-the-art MRI with quantified placental pathological outcomes in the primate for the first time. The final novelty of our study is the implementation of partial distance-based Redundancy Analysis - effectively removing animal-specific variance and revealing latent dimensions of pathology that would have otherwise remained obscure. Crucially, these findings were all demonstrated in healthy primate pregnancy - highlighting the spectral nature of pathological manifestation in relation to placental perfusion and fetal outcome.

In shifting away from qualitative placental assessment towards exhaustive, digital annotation, we provide a new framework for understanding placental pathogenesis in healthy and unhealthy pregnancy alike. Of note were several findings warranting further investigation: 1) the agreement of observed pathologies with historical clinical data; 2) the distinction and local co-expression of common MVM pathologies; 3) the connection between stromal mineralization, maternal perfusion, and fetal outcome; and 4) healthy pathological features observed in well-perfused cotyledons with chronically low blood flow.

### Primate Placental Pathology Quantification Aligns with Clinical Observations

The global pathological trends emerging from our healthy pregnancy cohort generally aligned with those observed clinically. Within the cotyledon, most injury manifested near the trophoblastic shell, in agreement with observations in human placentae [2]. Fibrin deposition, intervillous gaps, and stromal mineralization were the pathologies most frequently observed at term, all of which are known to commonly increase throughout gestation [36,43,44]. Simultaneously, inflammatory and non-inflammatory villous agglutination constituted the smallest proportion of total pathology, broadly indicative of lesions less common clinically [6,35]. We found agglutination pathologies most often occurred along the parenchymal border, whereas fibrin deposition and stromal mineralization most often occurred within the central parenchyma. This too aligns with prior observation; pathological and sub-pathological agglutination features are commonly observed at the maternal floor and periphery [35,38], whereas stromal mineralization and fibrin deposition often occur across the entire villous region [5,36,45].

### MVM-Related Pathologies are not associated with Chronic Malperfusion

Pathologies of MVM are categorized by their assumed connection to altered maternal blood perfusion at the placenta [38,46,47]. Despite its eponym, little work has directly connected quantified maternal blood perfusion with placental pathology manifestation. The prominence of MVM-associated Components 1 and 2 aligns well with prior observation, as these pathologies constitute the most common group of placental lesions and demonstrate high co-expression [7]. Surprisingly, we found across multiple modalities that these MVM-associated pathologies have little association with chronic hypoperfusion. As our methodology only captures three timepoints across mid to late gestation, it is possible that acute bouts of transient ischemia from maternal spiral arteries result in the observed MVM pathologies, as this phenomenon has been observed to occur naturally in the primate [48], and long hypothesized as a mechanism of adverse pregnancy outcome [49]. Future investigation will focus on the effects of transient ischemia on placental health *in vivo* and *in vitro*, to determine whether acute ischemia/reperfusion events manifest as MVM injury.

### Stromal Mineralization at Term is associated with Compensatory Perfusion

Multiple analysis modalities revealed that most distinct from MVM-associated injuries was stromal mineralization, the singular annotated pathology highly associated with FVM [36]. Furthermore, stromal mineralization demonstrated the strongest ties to local maternal blood perfusion, as increases were associated with poor perfusion and longitudinal compensation, while decreases of were associated with physiological fibrin and low flow cotyledons demonstrating adequate perfusion. The precise mechanism behind this mineralization is unknown, however, it has been hypothesized to occur by continued diffusion of minerals across the syncytia of pathological fetal villi with diminished flow [36]. Previously, Roberts et al. described increased chorionic mineralization within compensatory cotyledons of severely injured placentae [50]. Additionally, the connection between fetal villous damage and altered maternal hemodynamics at the placenta has been previously established in adverse outcome pregnancy [51,52]. The connection described here, between stromal mineralization, hypoperfusion, and a maternal hemodynamic response is novel observation in healthy primate pregnancy; demonstrating the significant, spectral relationship between pathological manifestation and perfusion locally.

### Physiological Fibrin Deposition and Healthy Placental Function

Finally, in Component 4 of our dbRDA analysis we show that reductions in stromal mineralization were associated with a subset of maternal floor fibrin – distinct from MVM-associated fibrin. We propose this fibrin represents Rohr’s and Nitabuch’s Fibrinoid, physiological fibrin theorized to protect villous tissue by reducing turbulent maternal blood flow while simultaneously providing a fetal buffer from maternal immune recognition/injury [26]. This Component 4 relationship was strongly associated with low cotyledon blood flow and volume across gestation, alongside sustained perfusion; in contrast with the high stromal mineralization cotyledons that demonstrated diminished perfusion and compensatory changes. It is likely that increased villous growth/volume of the cotyledon reflects inadequate maternal blood supply. Conversely, small cotyledons demonstrate little compensatory growth, alongside low blood flow and low FVM pathology due to their adequate level of perfusion. Together, these data work to disentangle optimal placental blood supply at a local vs global level - challenging the axiom that an increased blood flow parameter alone is indicative of healthier placentae and improved fetal outcome.

### Limitations and Future Directions

Our methods to quantify pathology at the placenta attempt to define fundamental pathological features; features that demonstrate extensive overlap across the many pathological categorizations described across decades of placental analysis literature [2,5,35]. As this was the first iteration, future increased magnification and specificity of histological annotation will provide ever-increasing resolution. For example, fibrin deposition was the sole pathology to not demonstrate a strong, positive association with its own occurrence across sub-tissues. The consolidation of many fibrin subtypes (physiological vs pathological, matrix-type vs fibrin-type, intra- vs intervillous [24]) represents an inherent limitation of this analysis, and likely contributed to obfuscation of intra-pathological, inter-pathological, and pathology-perfusion relationships. Future efforts will be geared towards delineating discrete fibrin manifestations. Additionally, the relatively low number of pregnancies and associated pseudoreplication limits the statistical validity and generalizability of the association between stromal mineralization and decreased fetal weight, and therefore should be interpreted cautiously. Future efforts will be made towards increasing the rhesus cohort size, pathological variance and including human placental tissue, to ensure these findings are both better validated and map onto real human placental physiology.

## Conclusions

Multiple groups have observed the need for a more objective assessment of baseline pathology at the placenta [2,53,54]. While unsupervised computational approaches have begun to fill this gap, it is our position that the best way to distill the knowledge gained from the past century of placental pathology investigation with current advanced analysis techniques is through applying digital expert-led annotation of placental injury and architecture alongside these novel technologies. Across pathological literature, digital annotations improve attention alignment in WSI classifiers [55], improve region labeling, and reduce noise [56]. Within placental pathology, annotation-implemented AI pipelines have already enabled structured mapping of placental cell-to-tissue architecture and WSI-based identification of preeclamptic placentae [57,58]. These studies support manual annotation of placental compartments and lesions as a practical way to steer models toward biologically meaningful regions, improve downstream adaptation, and provide a foundation for more grounded interpretability. In conclusion, we challenge the view of placental lesions as binary indicators of disease; through pixel-level annotation paired with longitudinal perfusion data we demonstrate that the boundary between health placental maturation and early insufficiency is graded and resolvable at the cotyledon level.

## ACKNOWLEDGEMENTS

The work was supported by the National Institutes of Health Grants R01-HD103443 and P51-OD011106 (to the Wisconsin National Primate Research Center [WNPRC]). JV was supported by NIH grants K01AI82448 and L70HD119811. A.K.S was supported by NIH grants AI175753-01 and HL163623-01A1. T.J.K was supported by the Brain & Behavior Research Foundation (National Alliance for Research on Schizophrenia and Depression; NARSAD) Young Investigator Award and the MQ Mental Health Research Fellowship. The content of this manuscript is solely the responsibility of the authors and does not represent the official views of the NIH. We gratefully acknowledge GE Healthcare for research support of University of Wisconsin-Madison, and AMAG Pharmaceuticals for providing ferumoxytol used for this study. We also thank the WNPRC Veterinary, Scientific Protocol Implementation, and Animal Services staff for providing animal care, and assisting in procedures including breeding, pregnancy monitoring, and sample collection.

