## Supplemental Figures and Tables for "Bridging Cotyledon Pathology and Perfusion in Healthy Primate Pregnancy"

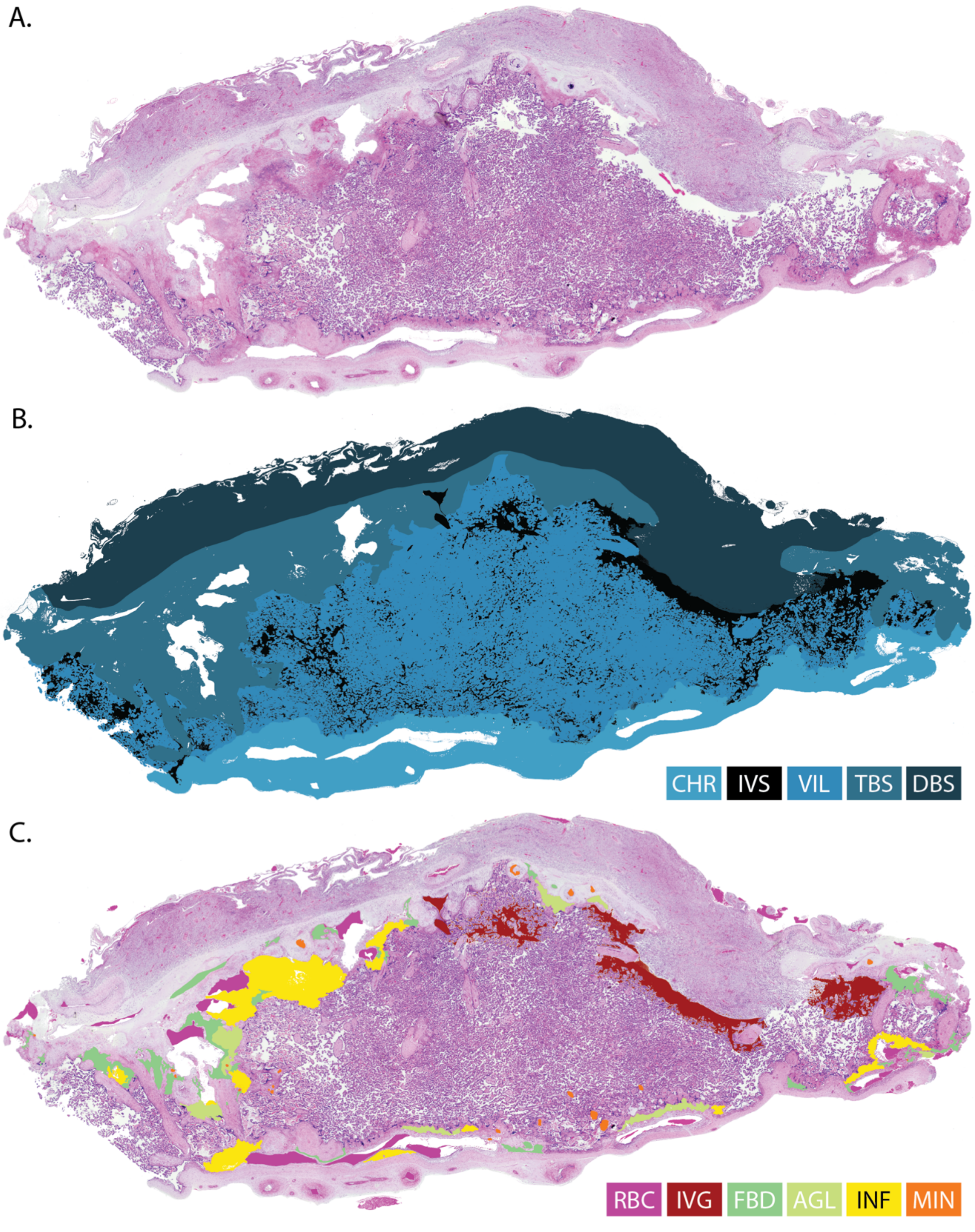


**<u>Supplemental Figure 1</u>. Representative images of stages of placental annotation.** A) An H&E-stained placental cotyledon scanned at 4x magnification. B) Sub-Tissue annotations: chorionic plate (CHR), intervillous space (IVS), placental villi (VIL), trophoblastic shell (TBS), and decidua basalis (DBS). C) Red blood cell (RBC) and pathology annotations: intervillous gaps (IVG), fibrin deposition (FBD), villous agglutination (AGL), inflammatory agglutination (INF) and stromal mineralization (MIN).

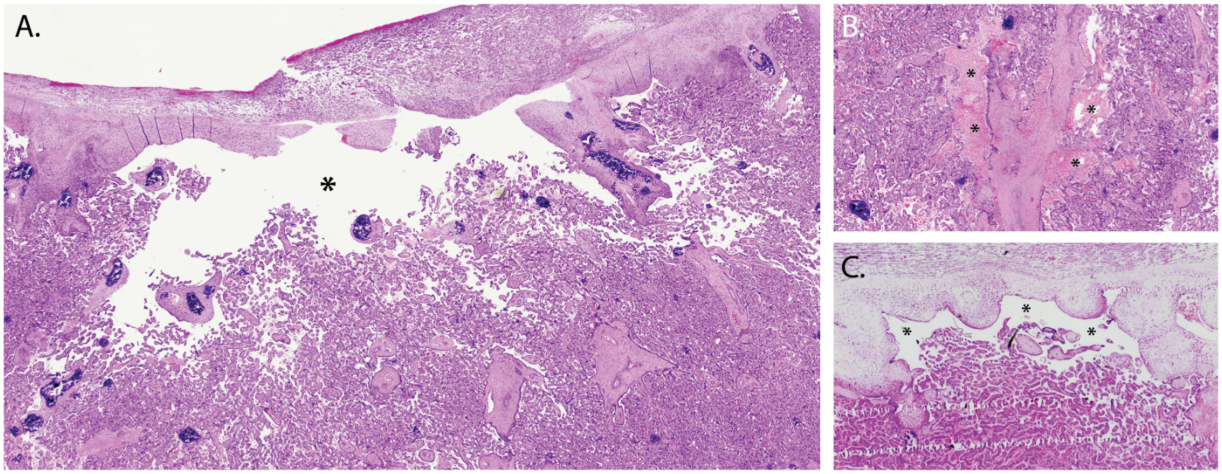


**<u>Supplemental Figure 2.</u> Representative images of cotyledon intervillous gaps.** A) A large gap in the intervillous space along the trophoblastic shell. B) Two areas of blood pooling in the central parenchyma, alongside a stem villus. C) A smaller gap in the intervillous space, along the trophoblastic shell. Asterisks "*" represent pathological areas of interest. All images taken at 4x magnification.

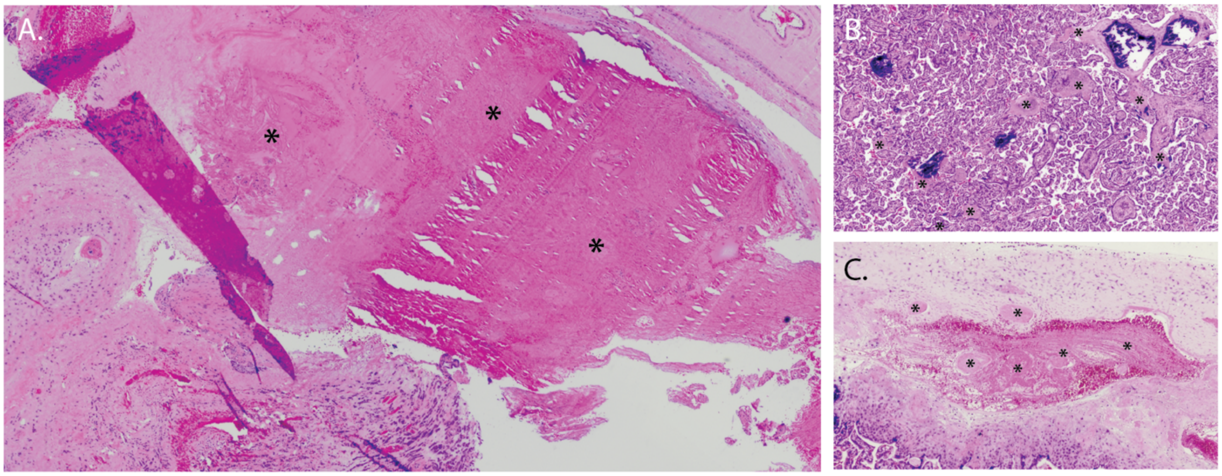


**<u>Supplemental Figure 3.</u> Representative images of fibrin deposition (FBD).** A) An area of mostly homogenous fibrin deposition along a placental septum. B) Diffuse FBD throughout the villous region. C) A pocket of partially coagulated blood within the trophoblastic shell, with FBD islands. Asterisks "*" represent pathological areas of interest. All images taken at 4x magnification.

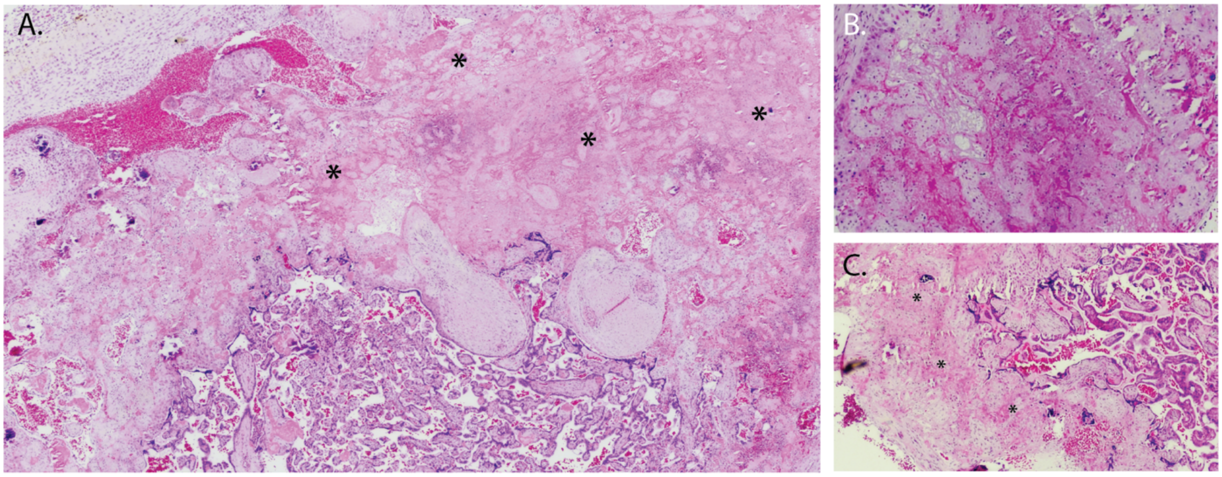


**<u>Supplemental Figure 4</u>. Representative images of villous agglutination (AGL).** A) AGL along the trophoblastic shell, with small areas of heavy leukocytic infiltrate throughout. B) AGL found within the central parenchyma. C) AGL along the chorionic plate. Asterisks "*" represent pathological areas of interest. All images taken at 4x magnification.

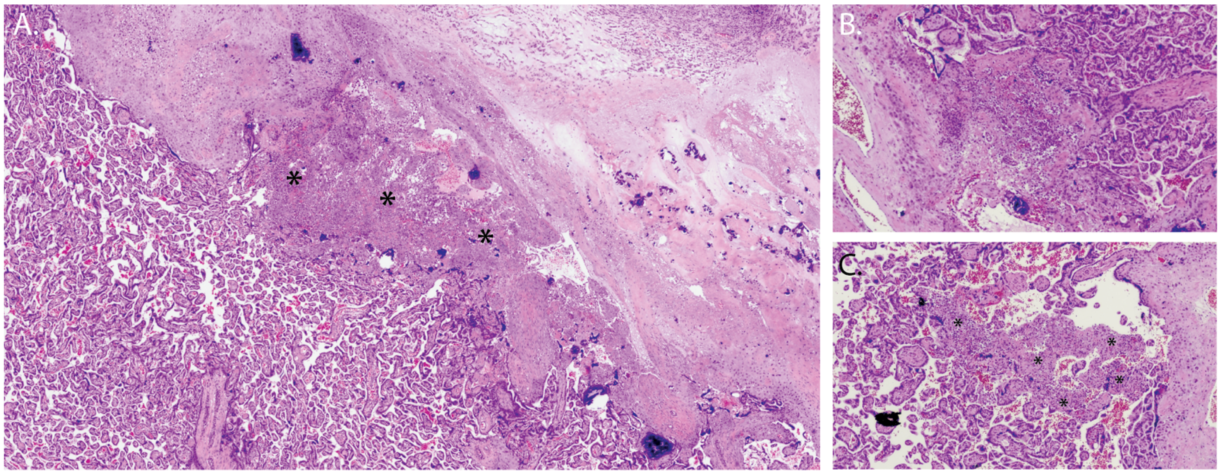


**<u>Supplemental Figure 5</u>. Representative images of inflammatory villous agglutination (INF)**. A) INF along the trophoblastic shell, with small areas of heavy leukocytic infiltrate throughout. B) INF within the central parenchyma. C) INF along the chorionic plate. Asterisks "*" represent pathological areas of interest. All images taken at 4x magnification.

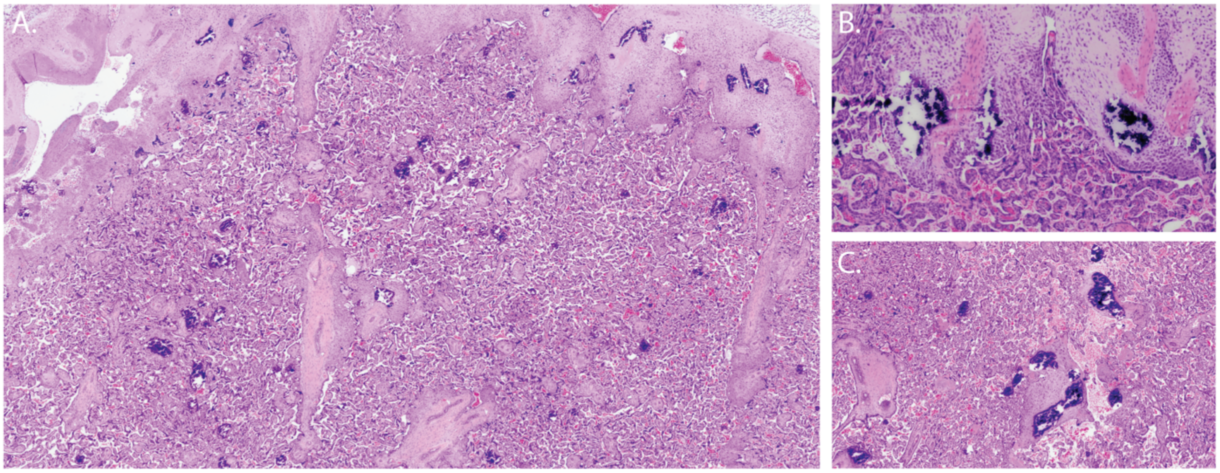


<u>**Supplemental Figure 6**</u>**. Representative images of stromal mineralization (MIN).** A) MIN across the villous region and trophoblastic shell. B) MIN within the stroma of anchoring villi. C) MIN within stem and floating villi. All images taken at 4x magnification.

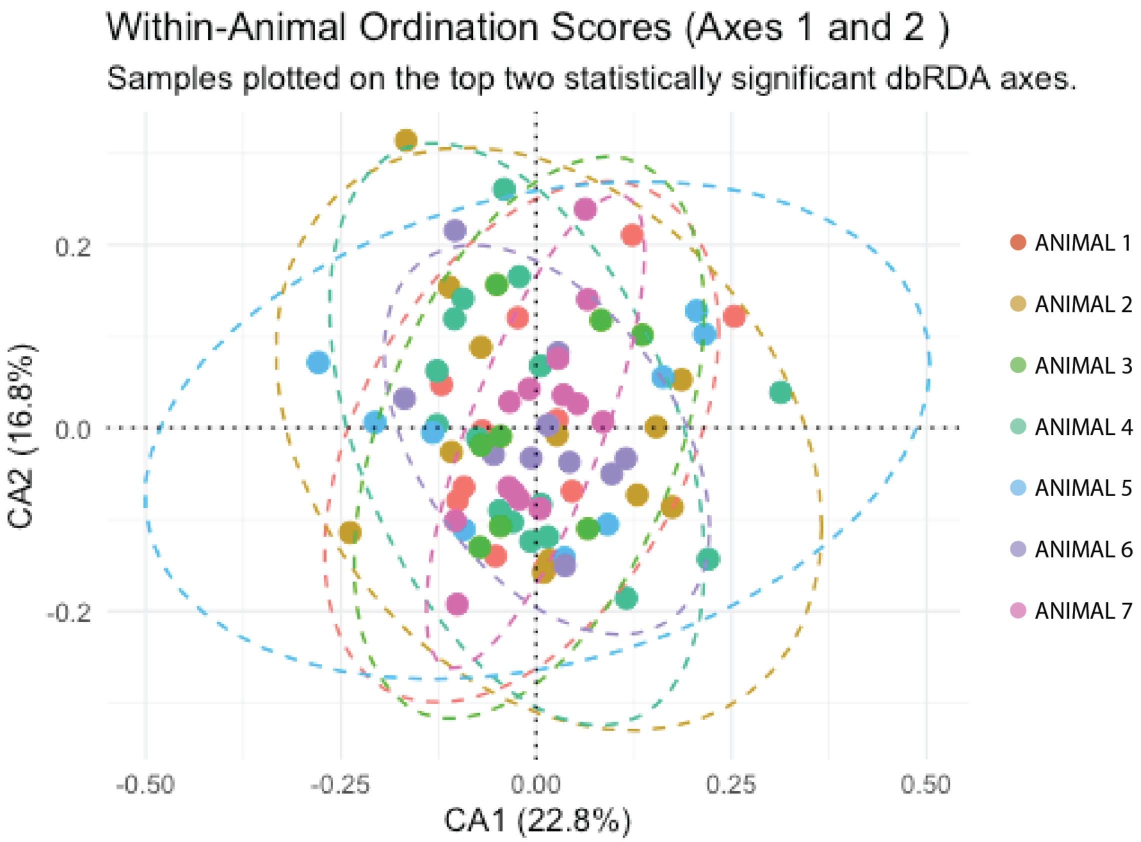


**<u>Supplemental Figure 7</u>. Removal of Animal Variance from Pathological Component Data.** Dot plot depicting the first two ordination axes from dimensionality reduction (CA1 and CA2) and their explained variances. Each dot represents individual cotyledon scores colored by animal. Uniform (random) distribution around the origin suggests the removal of animal-specific sources of variance was successful.

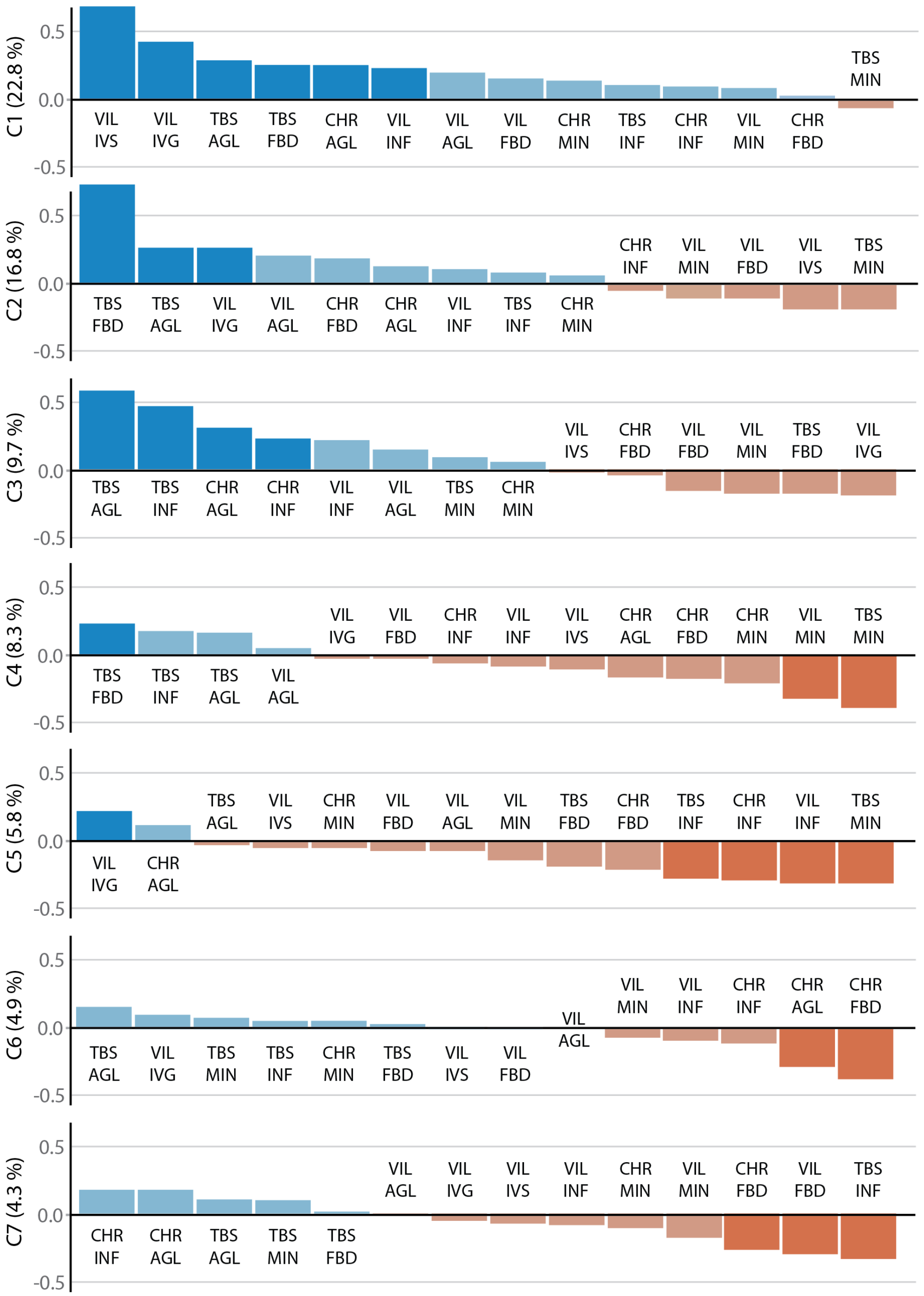


**<u>Supplemental Figure 8.</u> Pathological Components.** All pathological components demonstrated to significantly account for dataset variance, from most (top) to least (bottom) total variance accounted for. Y axis represents Spearman correlation values. Darker colored bars represent individual pathologies positively (blue) or negatively (orange) contributing to a given component significantly ($p<0.05$).

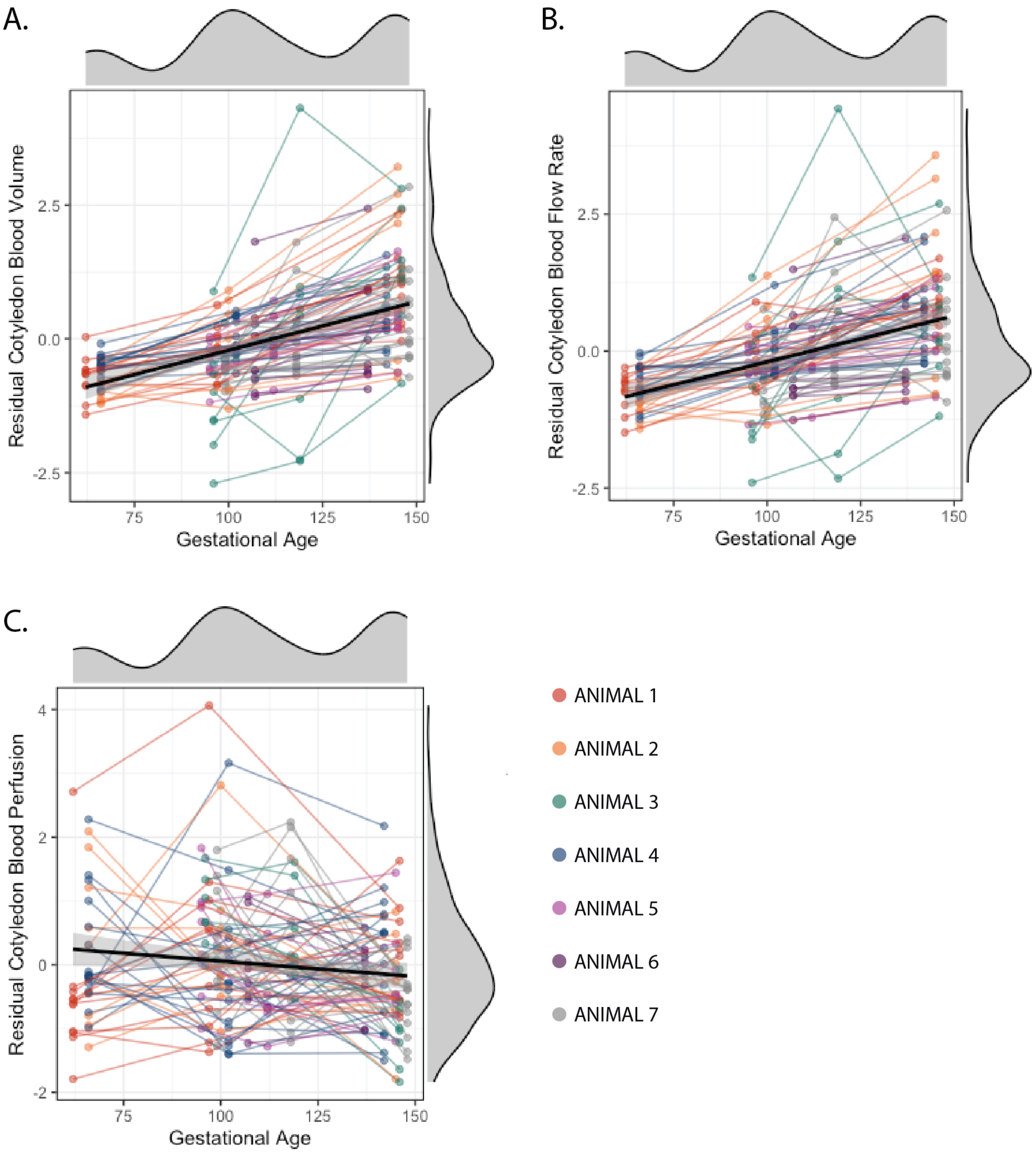


<u>Supplemental Figure 9.</u> **Mixed Model Analysis of Gestational Age and MRI Metrics.** Dot plots depict datapoints from all A) blood volume, B) flow, and C) perfusion MRI timepoints, from each cotyledon, in relation to gestational age. All relationships were found to be significant pre-correction, with blood volume and flow remaining significant following FDR correction.

**Supplemental Table 1**. Cotyledon sub-tissue annotation criteria.

| Cotyledon Sub-Tissue | Inclusion Criteria | Exclusion Criteria |
|---|---|---|
| Decidua Basalis (DBS) | Differs in cellular structure, frequency of nuclei, and increased hematoxylin staining compared to the TS. | Blood pooling outside, within, or in between shell and decidual layers. |
| Trophoblastic Shell (TBS) | Differs in structure and staining (lighter, usually more eosinophilic) between decidua and villi. TBS nuclei were generally more sparse. Anchoring villous tissue and agglutinated villi continuous with the TBS were included in TBS (no tissue break or dramatic staining color change). Where the TBS and CHR meet, distinctions were made by tissue structure and staining. When distinctions were not evident, the area was split half-way between most easily identifiable TBS and CHR areas. | Blood pooling between identifiable shell and villous regions, as well as pooling between shell and decidual regions. |
| Placental Villi (VIL) | All tissue that was not decidua, TS, or chorionic plate. Any blood pooling between shell and villi, or between chorion and villi, was considered placental villous area. | Any peripheral red blood cells and cellular/tissue debris. |
| Chorionic Plate (CHR) | Differs in structure and staining from villi. Chorionic stem villi emerging from the chorion or agglutinated tissue continuous with the chorion was included in chorionic plate area (no tissue break or dramatic staining color change). See above for criteria when the CHR and TBS met. | Blood pooling between chorionic and villous regions. |
| Intervillous Space (IVS) | All negative space, along with defined blood pooled RBCs between the placental villi. | Negative space and blood pooling within stem villi, negative space within diffuse, peripheral placental villi, negative space within villi near large tears/sectioning artifacts |

**Supplemental Table 2**. Cotyledon pathology annotation criteria.

| Pathology | Inclusion Criteria | Exclusion Criteria | Related Terms |
|---|---|---|---|
| Intervillous Gaps (IVG) | Gaps within the intervillous space located from the central villous region to the trophoblastic shell that were larger than 100,000 pixels. Diffuse and semi-structured red blood cells were included. | Large pockets of fibrin, diffuse RBCs, or gaps near histological tears/artifacts | Early intervillous thrombi, distal villous hypoplasia, loss of villous tissue |
| Fibrin Deposition (FBD) | Eosinophilic material, either a nucleated (fibrin-type) or sparsely nucleated (matrix-type). Only substantial, easily discernable regions were quantified (at least ~2,000 px large). May include lighter, land-locked eosinophilic villi surrounded by fibrin. If fibrin and RBCs were diffusely intermixed, regions composed of more than half fibrin deposition by area were included. | Large regions of light/moderate, eosinophilic tissue with sparse or moderate nuclei staining within the trophoblastic shell or chorionic plate | <u>Physiologic</u>: Rohr's/ Nitabuch's/Langhans' stria/fibrinoid<br><u>Pathologic</u>: Fibrinoid deposition, intravillous fibrin deposition, intervillous fibrin deposition, (massive) perivillous fibrin deposition, intervillous thrombi |
| Villous Agglutination (AGL) | Loss of villous structure. Intra- and perivillous fibrin present, potentially accompanied by diffuse RBCs. Tissue usually presents more eosinophilic but may have hematoxylin present. Agglutinated regions continuous with the TS or chorionic plate were included in those respective tissues. | Agglutinated regions with heavy dark dots (leukocytes) | Villous infarction, maternal floor infarction, perivillous fibrin deposition |
| Inflammatory Agglutination (INF) | Loss of villous structure with leukocyte infiltrate. Intra- and pervillous fibrin present, potentially accompanied by diffuse RBCs. Tissue presents as a mix of deep eosinophilic/hematoxylin with dark dots heavily present. Small ghost villi surrounded by inflammatory damage were also included. Agglutinated regions continuous with the TS shell or chorionic plate were included in those tissues. | Agglutinated regions with few or no areas of punctate hematoxylin stain (leukocytes). | Villous infarction, maternal floor infarction, perivillous fibrin deposition, chronic villitis |
| Stromal Mineralization (MIN) | Inclusion: Semi-circular, cracked hematoxylin-stained tissue, often with an empty center, confined to the villous stroma. | Syncytial calcification throughout villi and along the chorionic/TS border. | Calcification, segmented villous mineralization |
